# The brain prioritizes information that strengthens the structure of knowledge

**DOI:** 10.64898/2026.07.13.738178

**Authors:** Refael Tikochinski, India Pinhorn, Tal Nahari, Amber Duettmann, Tali Sharot

## Abstract

Humans learn surprisingly well from remarkably little. One way by which such feat may be accomplished is strategic information processing. In particular, the brain may prioritize new inputs that are likely to support efficient mental models. Here we show that people prefer semantic information that strengthens the small-world configuration of a knowledge system, a topology previously associated with creativity and comprehension. Such information is prioritized by the brain’s language network and elicits enhanced activation in the ventromedial prefrontal cortex, which is central to the brain’s valuation system. These findings (replicated across studies) reveal a principle of information prioritization: the brain assigns value and resources to knowledge according to its capacity to improve the structure of what we know.

## Introduction

Humans rely on internal models of the world to predict outcomes and generate novel solutions. These models incorporate semantic knowledge that captures the rules, relationships and causal structure of the environment^1–6^. Yet only a minute fraction of available information can be encoded: time, resources and computational constraints sharply limit what learners can process.^7^ Despite this, human cognition supports remarkable comprehension,^8,9^ abstraction^6^ and innovation,^10^ suggesting that while impoverished these models are remarkably effective.^9,11^

One way to accomplish this feat is by sampling information strategically.^12–17^ The brain may prioritize inputs that support compact, efficient mental models rather than processing information indiscriminately. Whether neural responses to semantic information follow such a principle, and the precise form that principle takes, is unknown.

Here, we test the hypothesis that the brain preferentially responds to information that reinforces a small-world knowledge structure. A small-world structure is an organizational principle that characterizes many systems from biological to man-made (such as social networks, transportation systems and neural networks).^15–18^ It is characterized by dense local clusters that connect nearby nodes and direct links between distant clusters. This organization is thought to reflect the relative optimization of local connections that create coherence and global connections across clusters which together facilitate a network’s function.^18–20^

Human semantic knowledge is often conceptualized as a network of interconnected concept^21,22^, where meaning is derived from the relationships among them^23,24^. Importantly, semantic knowledge also exhibit a small world topology^1,2,25–27^, balancing local coherence with access to distant associations. This structure has been linked to learning^1,26^ and creative thought,^2,28,29^ because dense local clustering supports categorization and generalization^1,3^ while short paths between distant domains facilitate creativity and innovation by enabling associations across otherwise remote concepts^3,28–30^.

Thus, the prediction is that people will exhibit a preference to engage with information that creates links between concepts that in turn strengthens the small-world structure of knowledge. This prioritization should be evident both in behavior and in neural responses to new information. To test this, we conducted three studies, including a behavioral experiment, a large language model validation, a primary functional magnetic resonance imaging (fMRI) experiment and an additional fMRI replication. We presented factual sentences to participants and examined whether they preferred those that reinforced the small-world structure of knowledge, and if neural responses showed prioritization to such stimuli.

In particular, we focus on two neural systems. First, as the information is presented as semantic text, we would expect neural prioritization within the language system to information that increases the small-world structure of knowledge. Key regions in this network include the left frontal lobe (in particular the inferior frontal gyrus), the left temporal lobe (in particular the posterior superior temporal gyrus and the middle temporal gyrus), the anterior temporal lobe, and the inferior parietal lobe (in particular the angular gyrus and supramarginal gyrus).^31–34^ Second, it has been shown that the brain signals the value of information using the same neural system and code as for primary rewards (e.g., food and water) and secondary rewards (e.g., material things).^35–39^ This is consistent with the ‘neural common currency’ hypothesis,^40–42^ which suggests that the brain uses a single, common scale to represent the subjective value of diverse options. This allows agents to compare qualitatively different alternatives to efficiently guide choice. The value of semantic information in particular has been shown to be represented in the ventromedial prefrontal cortex (vmPFC),^43^ which is the central node of the value system.^40^ We would thus expect the vmPFC to respond to the ability of information to increase the small-world configuration of a knowledge network if such characteristic is indeed valuable. Similarly, we will expect to see behavioral indices of this value.

If humans preferentially engage with information that reinforces the small-world structure of knowledge, this could be key in contributing to the effective organization of human knowledge. It would provide a mechanism that may help explain the surprising human ability to learn so much from so little.

## Results

To test the above hypothesis, we presented factual sentences to participants and asked them to indicate how much they liked each sentence and how much they wanted to engage with more sentences like it. We then examined whether these behavioral indicis of preference were associated with the impact of the sentence on the small world configuration of a knowledge network.

To do this we first had to build a knowledge network (Figure 1). We included 500 concepts in the network, which included all 169 concepts that were presented to participants in the factual sentences plus 331 concepts sampled from Wikipedia (see Methods for the sampling procedure). Using different knowledge systems with 250, 1000, and 2000 total concepts does not alter the results (see supplementary Table S1).

**Figure 1.**
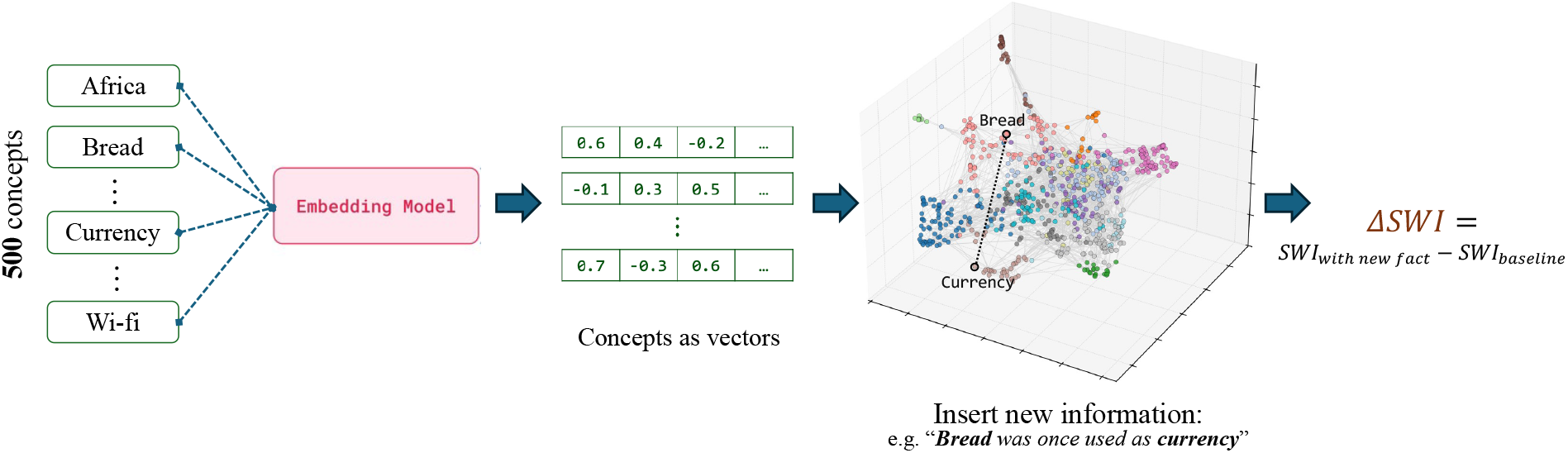
**Constructing a graph representation of a knowledge network.** We sampled 500 Wikipedia concepts, which included all the concepts in the stimuli presented to participants, plus 331 concepts selected using TF–IDF scores (see method for details). Using larger networks with more concepts provide the same results (see supplementary material). Concepts were embedded into a high-dimensional semantic space using the *wikipedia-en-embeddings* retrieved from huggingface.com. Pairwise cosine similarities between concept embeddings were used to construct a weighted graph (nodes = concepts; edges = cosine similarity if it exceeds the threshold; otherwise, 0) representing an approximation of a large-scale knowledge network. To quantify the impact of each piece of new information on the network, we calculated the Small-World Index (SWI) of the network before the new fact was added (SWI_baseline_) and after a piece of information was introduced (SWI_with_new_fact_) - that is, after strengthening/creating the connection between the two key concepts in the sentence - and calculated the difference between the two (ΔSWI= SWI_with_new_fact_ - SWI_baseline)_. This procedure was repeated for each of the 100 factual sentences, yielding a unique ΔSWI value for each fact.

The Wikipedia entry of each of the concepts was transformed into an embedding vector using the Open AI’s embedding model *“text-embedding-ada-002”* (retrieved from https://huggingface.co/datasets/Supabase/wikipedia-en-embeddings). Representing each concept via its full Wikipedia entry, rather than the embedding of the concept word alone captures its broader meaning, as large language models are context-dependent and a richer contextual basis yields a more comprehensive concept representation.^44,45^ Those embedding where then transformed into a graph space by calculating pairwise cosine similarities between all concepts to construct an undirected weighted graph where each node corresponds to a concept, and an edge between nodes exists if their cosine similarity is greater than a predefined threshold. Cosine similarity of Wikipedia entries reveals a relationship between concepts, not merely if they are alike. For example, the cosine between *teacher* and *school* is high despite the entries not representing the same concept. The threshold for the existence of an edge was set at the 80th percentile of the cosine similarity distribution, but the pattern of results remained unchanged when alternative thresholds (50th, 70th, or 90th percentiles) were applied (see supplementary Table S2). The weight of each existing edge is set to their cosine similarity value. We then computed the small-world index (SWI) of the resulting graph using the standard definition of SWI ^46^:

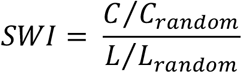

*C* is the clustering coefficient of the network, *L* is the average shortest path length, and *C_random_* and *L_random_* are the corresponding values for a random graph of equal size, density, and distribution of weights (see Methods for the weighted-graph formulations of *C*, *C_random_* , *L, and ^L^random* ^).^

To estimate the contribution of each new fact to the small-world structure of the network, we strengthened the edge weight between the nodes corresponding to the two key concepts in the factual sentence using the formula:

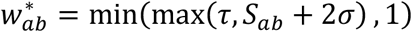

*S_ab_* is the original cosine similarity between concepts *A* and *B*, *σ* is the standard deviation of the cosine similarity distribution (see methods), and *τ* is the threshold defined for the graph. This ensured that the new weight is at least as high as the inclusion threshold and does not exceed the maximum allowable similarity value of 1. We then recalculated the small-world index of the updated graph, and defined the change in small-worldness due to the inclusion of a new fact as:

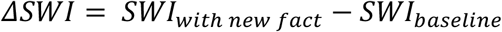

### Humans prefer information that enhances the small world configuration of the knowledge network

In **Experiment 1** ninety-three participants were presented with 100 facts (7s each). They indicated how much they liked each fact (on a scale from 1‘didn’t like it at all’ to 5‘liked it very much’; self-paced) and how much they wanted to consume more related facts (self-paced; on a scale from 1 ‘not interested at all’ to 5 ‘very interested’; Figure 2a). They also rated how surprising they found the fact to be (on scales from 1 ‘not surprised at all’ to 5 ‘very surprised’; self-paced). Participants’ average scores were: liking (Mean = 3.3, SD = 1.08), wanting (Mean = 3.31, SD = 1.11) surprising (Mean = 3.27, SD = 1.37).

**Figure 2:**
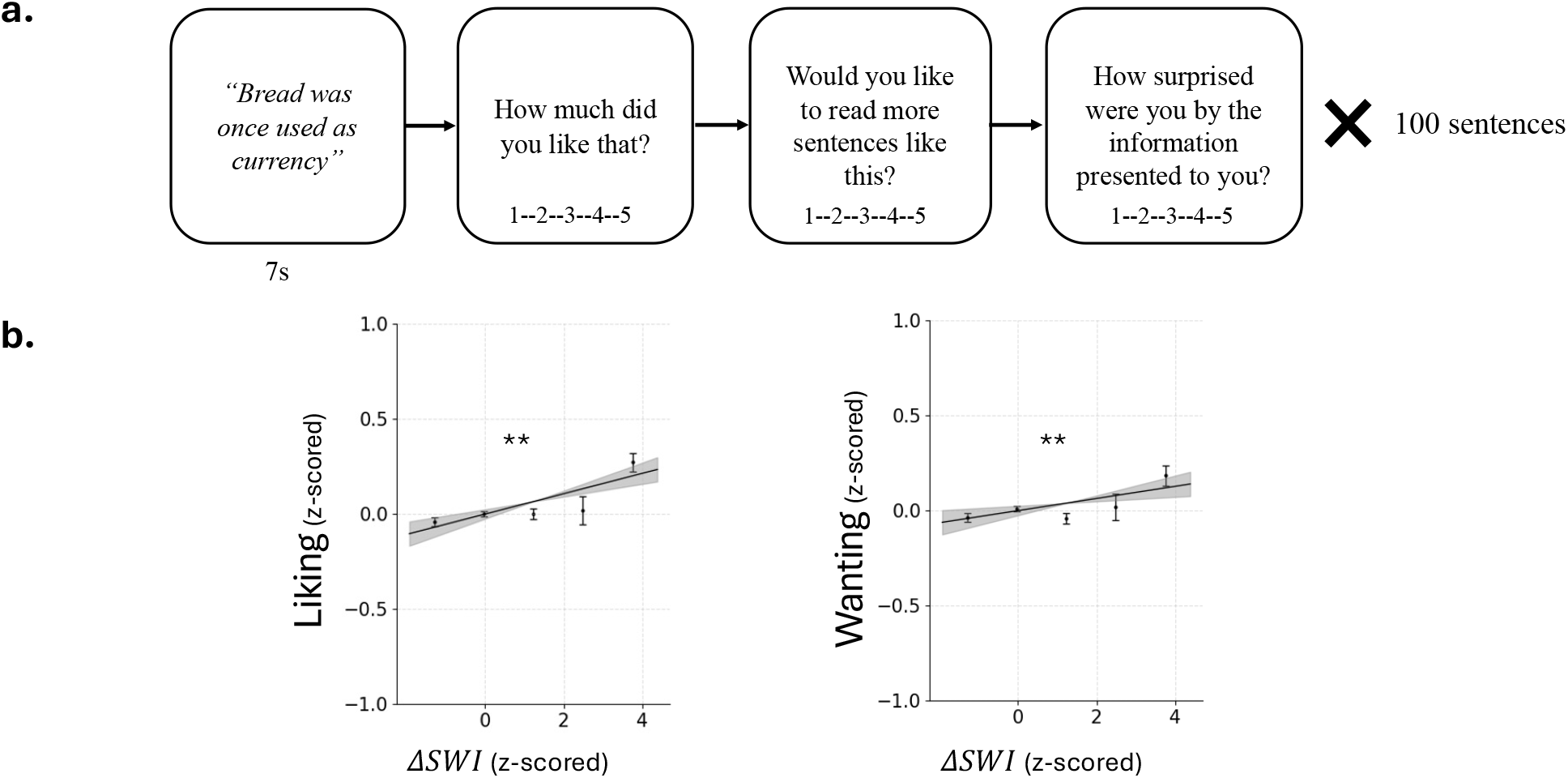
**Information that increases the small-world configuration of a knowledge system is preferred. (a). Task**. Participants were presented with 100 short factual sentences (7 sc) and indicted how much they liked the fact, how much they wanted to read more related facts, and how surprising they found it. **(b)** Facts that increased the small-world index (ΔSWI) were liked more and participants wanted to receive more related facts. This effect remained after controlling for surprisal and cosine similarity between the two concepts mentioned in the fact, as well as age, gender, and education. The line indicates the fixed effect estimated by the linear mixed-effects model fit to all (unbinned) data, and the shaded area represents the 95% confidence interval. For illustrations purposes only we also plot data binned into five equal-width intervals of the z-scored ΔSWI values: mean ± SEM; ** = p < .001.

Strikingly, we found that facts that increased the small-world configuration of the network (i.e., that produced larger *ΔSWI*) were liked more (linear mixed-effects model predicting liking from *ΔSWI* as fixed and random effects: β = 0.052, *SE* = 0.01, *t*(92) = 5.054, *p* = .0002; Figure 2b.). Moreover, participants indicated that they wanted to learn more facts related to those that increased the small-world configuration of the network (linear mixed-effects model predicting wanting from *ΔSWI* as fixed and random effects: β = 0.031, *SE* = 0.01, *t*(92) = 3.10, *p*=0.0025; Figure 2b). The same results hold when controlling for surprisal ratings and for the cosine similarity between the two concepts (both included as fixed effects and random slopes) as well as for participants’ age, gender, and education as fixed effects: (ΔSWI predicts *liking*: β = 0.048, *SE* = 0.01, *t*(96.25) = 4.688, *p*=0.0001; and *wanting*: β = 0.027, *SE* = 0.01, *t*(91.99) = 2.911, *p*=0.0045). Thus, participants are preferentially drawn to information that has the greatest potential to improve the small-world configuration of the knowledge system. The effects cannot be explained by how surprising the new fact is nor by whether the fact connects similar/dissimilar concepts, as these were controlled for in the model.

How much people liked and wanted information was associated with both the change to the clustering of the network (ΔC) and the change to the average path length of the network (ΔL). When examined separately, each component predicted liking and wanting (*liking*: ΔC: β = 0.023, *SE* = 0.02, *t*(91.98) = 1.99, *p*=0.049; ΔL: β = -0.046, *SE* = 0.01, *t*(91.99) = -4.662, *p*=0.0001; *wanting*: ΔC: β = 0.022, *SE* = 0.01, *t*(91.98) = 2.052, *p*=0.043; ΔL: β = -0.021, *SE* = 0.01, *t*(91.99) = -2.193, *p*=0.0308).

A model comparison using the Bayesian Information Criterion (BIC), which penalizes models for the number of parameters, favored the ΔSWI model over models containing only ΔC (liking: ΔBIC = 22.15; wanting: ΔBIC = 3.22) or only ΔL (liking: ΔBIC = 10.07; wanting: ΔBIC = 4.09). Moreover, a model comparison further revealed that the ΔSWI model also performed better than a model that included both ΔC and ΔL as fixed and random effects (liking: ΔBIC = 41.88; wanting: ΔBIC = 37.05). Together, these results indicate that the composite small-world index model explains participants’ attraction to information better than models that include only clustering and/or path length, suggesting that the attraction is driven by sensitivity to the joint optimization of local clustering and global path efficiency rather than to either structural property in isolation.

### LLMs indicate preference for information that enhances small-world configuration

We next examined whether Large Language Models also express preference for facts that increased the small-world configuration of a network. The rational was not that LLMs have preferences per-se, but that because they are trained on human data, they may reflect human preference. To that end we asked Open AI’s GPT-4o to rate all 100 facts on liking and wanting (see Method for prompt). We did this for 50 independent runs (‘virtual participants’). On each run each sentence was rated in isolation, without memory of previous trials (see Methods). Ratings were analyzed exactly as for human participants. The LLM showed the same preference as humans: ΔSWI significantly predicted both liking (without controls: β = 0.040, SE = 0.017, t(29.10) = 2.217, p = .0347; with controls: β = 0.036, SE = 0.017, t(29.45) = 2.10, p = .044) and wanting (without controls: β = 0.050, SE = 0.014, t(29.01) = 2.751, p = .0101; with controls: β = 0.056, SE = 0.013, t(29.46) = 3.069, p = .0045). The LLMs’ explicit preference provides a lens into latent patterns of human-generated text. In other words, the result suggests the preference is widespread and can be inferred from the human record.

### fMRI design

Thus far we see that information that increases the small-world configuration of a network is preferred. Next, we investigated whether this impact of information on a knowledge network is tracked by the human brain, particularly within neural systems associated with reward and language processing.

To this end, we recruited thirty-three new participants in **Experiment 2** who read the same 100 sentences from Experiment 1 while undergoing a functional MRI scan (Figure 3a). Each sentence was presented for 5 seconds and followed by a liking rating (‘How much did you like that?’ on a scale from 1 ‘did not like at all’ to 5 ‘like it very much’, self-paced). Each trial was preceded by a jittered fixation (3.5, 5.5, or 7.5 seconds in random order). After scanning, participants re-read all sentences and again provided liking ratings, which were correlated with the in-scan ratings (r = 0.49, p < 0.001). This post-scan procedure followed the same format as the behavioral task: participants indicated ‘How much did you like this fact?’ (from 1 ‘didn’t like it at all’ to 5 ‘liked it very much’, self-paced) and ‘How much do you want to learn more facts like this?’ (from 1 ‘not interested at all’ to 5 ‘very interested’). In addition, they indicated ‘How familiar was this information to you?’ (from 1 ‘not familiar at all’ to 5 ‘very familiar’).

**Figure 3.**
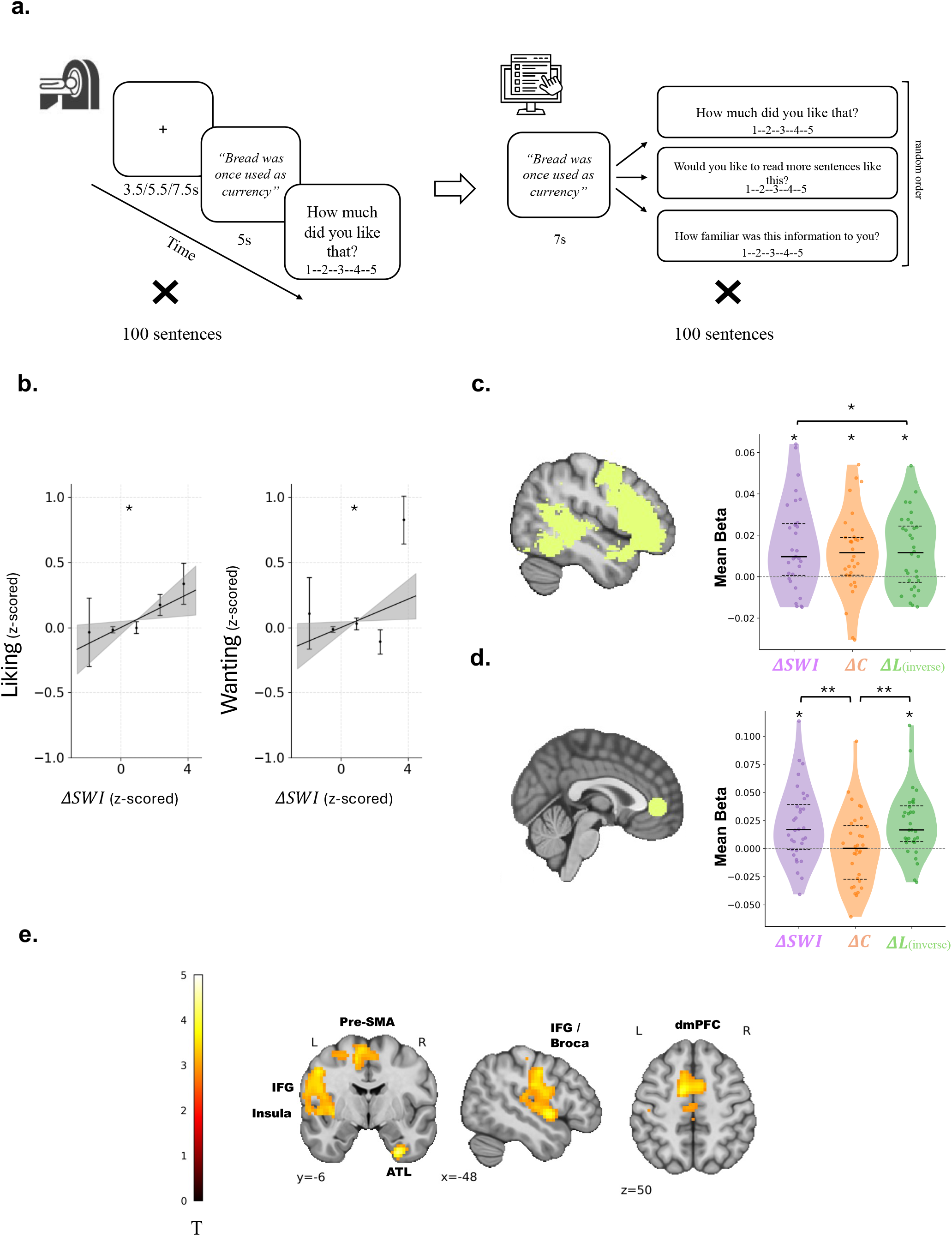
Neural prioritization of information that increases the small-world configuration of a knowledge network. (a) Task. Participants (n=33) read 100 factual sentences during fMRI scanning. Each sentence was presented for 5 seconds, followed by a liking rating (1–5 scale) and preceded by a jittered fixation. Following the scan, outside the scanner, participants read the sentences again and rated them on ‘liking’, ‘wanting’, and ‘familiarity’ using 1–5 scales. The order of the sentences and the questions were randomized. **(b)** Facts that increased the small-world index (ΔSWI) were liked and wanted more by participants. This effect remained after controlling for familiarity and cosine similarity between the two concepts mentioned in the facts, as well as age, gender, and education. The line indicates the fixed effect estimated by the linear mixed-effects model fit to all (unbinned) data, and the shaded area represents the 95% confidence interval. For illustrations purposes we also plot data binned into five equal-width intervals of the z-scored ΔSWI values: mean ± SEM**. (c)** *(Left)* Region of interest (ROI) derived from a Neurosynth meta-analytic map for the term ‘language’ (1,101 studies; FDR q < 0.01), encompassing voxels predominantly in the bilateral superior temporal gyrus (STG) and left inferior frontal lobe. *(Right)* Averaging all voxels within this language ROI showed that BOLD response was parametrically modulated by the impact of new information on the small-world index (ΔSWI) of the knowledge network as well as on the change in clustering coefficient (ΔC) and change in average path length (ΔL) of the network. For illustration purposes, we inverted ΔL in the figure so that all effects were displayed in the same direction, facilitating comparison of effect magnitudes across network measures. Statistical comparisons between ΔL and ΔSWI/ΔC were also conducted after inverting the scale of ΔL. **(d)** *(Left)* vmPFC ROI defined as a 10-mm sphere around the peak reward-association identified via Neurosynth (MNI: 0, 44, 0; Z = 7.31). *(Right)* Averaging all voxels within this vMPFC ROI showed that BOLD response was parametrically modulated by the impact of new information on the small-world index (ΔSWI) of the knowledge network as well as change in average path length (ΔL) of the network but not by the change in clustering coefficient (ΔC), suggesting that vmPFC valuation responses were primarily driven by the integrative, path-shortening properties of incoming information. As in (c), the ΔL axis was inverted for visualization purposes. **(e)** Exploratory whole brain analysis showing regions where activity is positively parametrically modulated by ΔSWI (p<.001 voxel-wise, cluster-corrected at FWE p<.05). Higher ΔSWI was associated with increased activity in medial prefrontal and superior frontal cortex (overlapping the medial multi demand network), a left-lateralized language network spanning inferior frontal and temporal regions, and bilateral cerebellum and ventral temporal cortex including the right anterior temporal lobe. In the violin plots (c & d), shaded areas show the probability density of the data at different values, smoothed by a kernel density estimator. Each dot represents a participant; larger dashed line represents the median; the two smaller dashed lines correspond to the first and third quartiles; ΔL beta estimates are sign-inverted for visualization. * = p<.05, ** = p<.01, n.s. = not significant.

Once again, we found that facts which increased the network’s small-world configuration were liked more (linear model exactly as in Experiment 1, without controls: β = 0.054, SE = 0.022, t(26.01) = 2.43, p = .021; with controls: β = 0.057, SE = 0.022, t(25.93) = 2.57, p = .016), and participants wanted to learn more related facts (without controls: β = 0.043, SE = 0.022, t(24.0) = 2.162, p = 0.040; with controls: β = 0.043, SE = 0.022, t(23.9) = 2.129, p = 0.043; Figure 3b).

### The language system prioritizes information that enhances the small-world configuration of knowledge

We first set out to examine if the language system is preferentially engaged when processing information that increases the small-world configuration of semantic knowledge. To that end, we defined a region of interest (ROI) map using neurosynth.org (note, that in Experiment 3 we replicate the results using ROIs which were defined functionally for each participant). Specifically, we obtained an automated meta-analytic association map for the term ‘language’ from Neurosynth.org, based on 1,101 studies. The map was thresholded at FDR q < 0.01 and reflects the relative selectivity with which individual voxels are associated with the term ‘language’, quantified by comparing reported activation frequencies in studies containing the term versus those that did not (see ref.^47^ for methodological details). The ROI that was generated included predominately voxels in the bilateral superior temporal gyrus (STG) and the left inferior frontal lobe (Figure 3c).

For each participant we then conducted a parametric analysis examining BOLD response at the time the participant viewed the stimuli (5s). Activity was modeled as a boxcar function and parametrically modulated by the ΔSWI of each stimulus. We controlled for variables of no-interest by adding then as additional parametric modulators in the model (these included familiarity ratings, cosine similarity between the two concepts in each sentence, number of words in the sentence, language model-based perplexity (a.k.a. ‘surprisal’), and serial position). Motion-related and physiological confounds were included as nuisance regressors (GLM 1, see Methods for details). For each participant we then calculated across all voxels in the ROI the mean beta coefficient for the parametric modulator of ΔSWI. This revealed a significant positive effect, indicating that the language network was preferentially engaged when processing information that increased the small-world structure of a knowledge network (mean Beta = 0.015, SE = 0.006, t(32) = 2.245, p = .017; Figure 3c).

Because a small world structure is characterized both by high clustering and short average path length, it is possible that the language network was preferentially responding to one of these features or both. To test this, we created a new GLM (GLM2) which was identical to the first, but instead of ΔSWI as the main modulator we entered the change to the clustering of the network when inserting the new fact (ΔC) and the change to the average path length of the network when inserting a new fact (ΔL) as the main modulators.

Averaging the beta of these parametric modulators across voxels in the language ROI revealed that BOLD signal was positively associated with both. Specifically greater BOLD response was observed during the presentation of facts that increase the clustering coefficient of the knowledge network (mean Beta: 0.01, SE= 0.005, t(32)= 2.109, p = .04) and decreased the average path length of the knowledge network (mean Beta: -0.014, SE= 0.005, t(32)= 2.439, p = 0.02; Figure 3c). Interestingly, the beta of the ΔSWI parametric modulator was greater than the beta of the ΔC (mean difference= 0.004, t(32)= 2.10, p = .04) or ΔL (mean difference after inverting the scale of the ΔL betas = 0.002, t(32) = 2.29, p = .028), suggesting that the language system was specifically prioritizing information that balanced clustering with a reduction of path length across the knowledge system.

The results thus far suggest that the language system prioritizes semantic information that support a configuration of knowledge previously associated with learning^1,26^ and creativity.^2,28,29^ Next, we examined if the brain’s core value region shows a similar pattern of activation.

### Core value region (the vmPFC) preferentially responds to information that supports the small-world configuration of knowledge

If a small world configuration promotes the function of a knowledge network, for example by facilitating creativity and understanding,^1,26^ then the brain’s reward system may be sensitive to this value. The vmPFC is thought to be the core brain region which signals the value of stimuli, whether it is food, money, art, mates or information.^37–39^ We thus set out to examine if the vmPFC tracks the impact of information on the small-world configuration of knowledge. To that end, we again used neurosynth, this time to identify a ROI within the vmPFC. In particular, we entered the term ‘reward’ into Neurosynth.org^47^ to detect the peak voxel in the vmPFC associated with this term [MNI: 0, 44, 0] (Z = 7.31) and defined a 10-mm-radius sphere around it.

For each participant, we tested whether BOLD responses across all voxels averaged within this ROI was associated with the ΔSWI parametric modulator from GLM1. Indeed, we found that activity in the vmPFC ROI was positively associated with ΔSWI (mean Beta: 0.022, SE= 0.01, t(32)= 2.224, p = .034; Figure 3d) implying that this valuation region is tuned to the structural impact of information on knowledge.

We next examined if this association was due to a response to the change in the clustering coefficient of the knowledge network and/or to a change in the average path length of the knowledge network. Using GLM2, we found that BOLD signal within the vmPFC ROI tracked the impact of information on average path length (mean β = -0.022, SE = 0.009, t(32) = -2.606, p = .014) and not its impact on the clustering coefficient (mean β = 0.004, t(32) = 0.046, p = .96; Figure 3d). This suggests that the vmPFC’s response to ΔSWI associated with information was primarily driven by the impact of information on reducing average path-length.

### Exploratory analysis shows neural prioritization within the multi-demand network of information that supports a small-world configuration of knowledge

Finally, we conducted an exploratory whole brain analysis using GLM1 to examine for additional brain regions that track the impact of information on SWI (voxel-wise *p* < .001 with cluster-level FWE correction at *p* < .05). This exploratory analysis revealed that the impact of information on SWI (ΔSWI) was positively correlated with BOLD response in medial component of the multiple-demand (MD) network^41^ including a large medial frontal cluster (14,849 mm³, peak: MNI [–9, 3, 44], t = 4.18), which extended dorsally into the superior frontal cortex (subpeaks: [–6, –9, 64], t = 3.71; [6, 0, 54], t = 3.53), and laterally into superior frontal regions ([–24, 0, 61], t = 3.49; Figure 3e, Supplementary Table S3).

Moreover, ΔSWI was associated with increased activity in a large left-lateralized language network (21,383 mm³, peak: [–60, 3, 18], t = 4.28), spanning left inferior frontal and premotor regions, extending to the anterior insula and temporal pole ([–48, 3, –2], t = 3.96), and inferior frontal regions including Broca’s area ([–51, –6, 41], t = 3.73), with additional activation in left postcentral regions ([–51, –12, 24], t = 3.68; Figure 3e, Supplementary Table S3).

Lastly, we observed activation in bilateral cerebellum and ventral temporal regions (16,037 mm³, peak: [9, –27, –32], t = 4.99), including the right anterior temporal lobe (vATL: [30, –3, –42], t = 4.42), bilateral cerebellum ([15, –42, –22], t = 3.95; [–6, –51, –19], t = 3.92), and parahippocampal/fusiform areas (Figure 3e, Supplementary Table S3). This cluster encompasses regions critical for semantic integration and predictive processing, with the ventral anterior temporal lobe serving as a semantic ‘hub’^50^. Past studies have also suggested that the parahippocampus facilitates retrieval of semantic associations from long-term memory^51^. These results in combination with the ROI findings suggest that information that increase the small-world configuration of knowledge preferentially engages both domain-general attentional control systems and domain-specific language regions.

### Replication of neural and behavioural findings (Experiment 3)

To assess the replicability of our findings, we repeated the behavioral and neural analyses on an independent dataset from our lab^43^. In that study, 31 participants underwent fMRI scanning while viewing 60 factual sentences (each for 4s) followed by a liking scale (self-paced; from 1=’dislike a lot’ to 5=’like a lot’). This was followed by a display confirming their rating (0.5s) and a jittered fixation cross (3.5-7.5s, mean 4s; see more details in ^43^). After scanning, participants re-read the sentences outside the scanner and again rated their liking on a 1–5 scale (r=0.71 between in-scanner and post-scan liking ratings). They also rated their familiarity with each sentence on a 1–5 scale. Additional rating measures were collected in the original study for purposes of that study (see ^43^ for details). 45 of those sentences shared the same two-concept structure used in the current study, namely, sentences that explicitly link two distinct concepts. Three of these sentences overlapped with those used in Experiments 1 and 2 and the rest were unique. We computed the ΔSWI for each sentence using a knowledge-network that included the 90 concepts drawn from these 45 sentences, supplemented by 410 additional concepts sampled using the same sampling procedure, yielding a total of 500 concepts.

A linear mixed-effects model confirmed that, as in Experiments 1 and 2, sentences that increased the ΔSWI were liked more (β = 0.069, SE = 0.027, t(28.04) = 2.567, p = .015; Figure 4a). This effect remained significant after controlling for cosine similarity between the two concepts, familiarity rating, age and gender (β = 0.059, SE = 0.029, t(28.97) = 2.04, p = .049) demonstrating that participants’ preference for information that enhances the small world configuration of knowledge generalizes to an independent sample and stimulus set.

**Figure 4.**
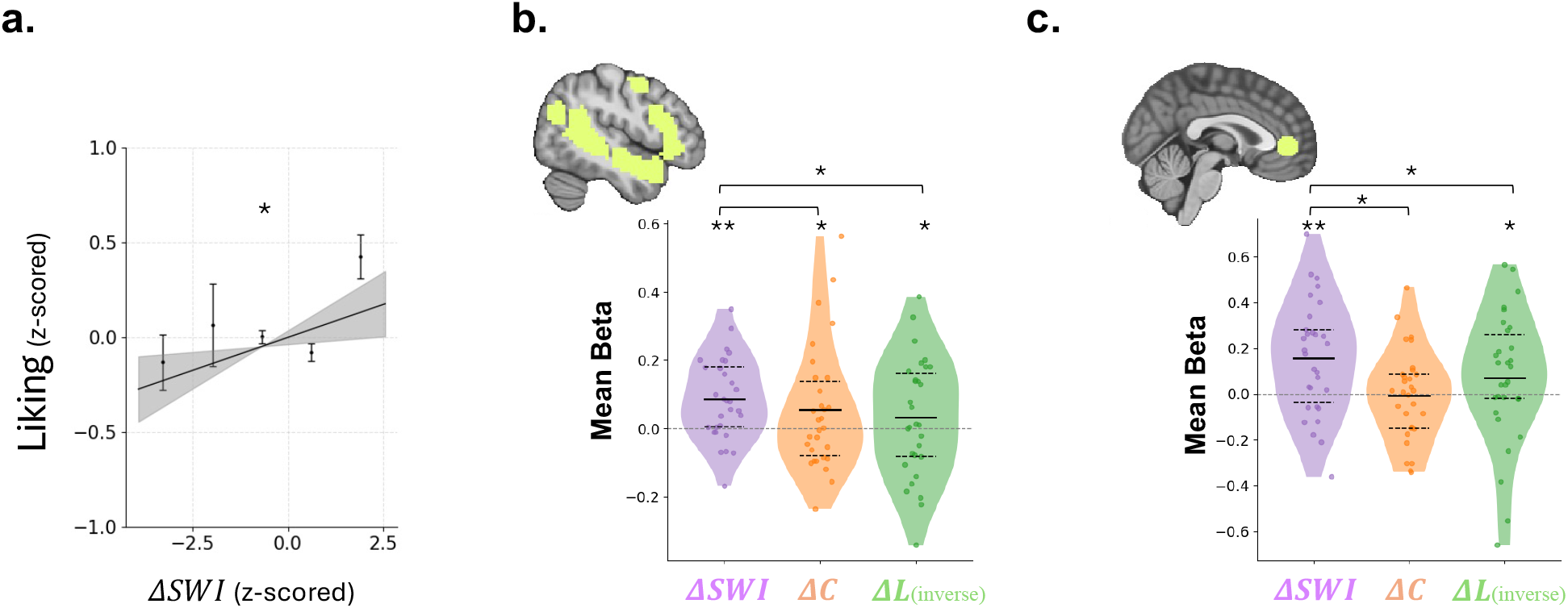
Replication and genelarization of behavioral and neural findings in an independent dataset (Experiment 3). **(a)** Facts that increased the small-world index (ΔSWI) were liked more by participants (n=31) in an independent sample. This effect remained significant after controlling for cosine similarity between the two concepts, familiarity, age, and gender. The line indicates the fixed effect estimated by the linear mixed-effects model fit to all (unbinned) data, and the shaded area represents the 95% confidence interval. For illustrations purposes we also plot data binned into five equal-width intervals of the z-scored ΔSWI: mean ± SEM **(b)** Averaging all voxels within the participant-specific functionally defined language ROI showed that BOLD response was parametrically modulated by the impact of new information on the small-world index (ΔSWI) of the knowledge network as well as on the change in clustering coefficient (ΔC) and change in average path length (ΔL) of the network. As in Figure 3(c&d), the ΔL axis was inverted for visualization purposes. **(c)** Averaging all voxels within the vmPFC ROI (defined as in Exp 2) showed that BOLD response was parametrically modulated by the impact of new information on the small-world index (ΔSWI) of the knowledge network as well as change in average path length (ΔL) of the network but not by the change in clustering coefficient (ΔC), replicating the valuation response pattern observed in Experiment 2. As in Figure 3(c&d), the ΔL axis was inverted for visualization purposes. * = p < .05, ** = p < .01, n.s. = not significant. In the violin plots (b & c), shaded areas show the probability density of the data at different values, smoothed by a kernel density estimator. Each dot represents a participant; the larger dashed line represents the median; the two smaller dashed lines correspond to the first and third quartiles.

One advantage of this dataset is that the scanning protocol included a standard language functional localizer task in which participants were presented with sentences and non-word strings^31^. None of the sentences in this localizer task overlapped with sentences presented in the main ‘liking’ task. Language-selective ROIs for each participant were identified by contrasting BOLD response while participants were presented with sentences versus when they were presented with non-word strings. The 10% of voxels showing the strongest contrast^43^ within the core language network derived from^52^ were defined as the ROI (note that this is exact method was used in^43^). We then used GLM1 to extract ΔSWI parametric beta estimates across all voxels within each participant’s idiosyncratic language ROI. We found that the language network showed greater activation for information that increased the small-world configuration of the knowledge network (mean ΔSWI beta = 0.072, SE = 0.026, t(30) = 2.788, p = .009; Figure 4b). Moreover, GLM2 revealed that the BOLD signal within participants’ idiosyncratic language ROIs tracked the impact of information on average path length (ΔL; mean beta = −0.025, SE = 0.022, *t*(30) = −2.05, *p* = .049) as well as on the clustering coefficient (ΔC; mean beta = 0.055, SE = 0.033, *t*(30) = 2.302, *p* = .028; Figure 4b).

Next, we examined the BOLD response in the vmPFC ROI, defined as in Experiment 2. Again, as in Experiment 2 we found that stimuli that increases the ΔSWI elicit greater BOLD response in the vmPFC (mean Beta: 0.158, SE= 0.046, t(30) = 3.445, p = .002; Figure 4b). Moreover, we found again that the BOLD signal in the vmPFC was also associated with the change to the average path length of the knowledge network (ΔL; using GLM2; Mean Beta = - 0.096, SE= 0.045, t(30) = - 2.022, p = .026) and not with the change in the clustering coefficient (ΔC; Mean Beta = -0.001, SE= 0.038, t(30) = -0.029, p = .97; Figure 4b) replicating the observation that the reward system’s sensitivity to ΔSWI is driven selectively by reductions in average path length. Together, the results of Experiment 3 replicate those of Experiment 2, showing neural prioritization within the vmPFC and the language network of information that increases the small world configuration of knowledge. This replication is not only on different participants, but also using different stimuli and a different knowledge network, which suggest generalization of the results.

A whole-brain analysis (*p* < .001 voxel-wise, cluster level FWE correction *p* < .05) revealed significant positive associations between ΔSWI and BOLD signal in left-lateralized language and frontoparietal regions. A large cluster was observed in the left supramarginal gyrus extending into the inferior parietal lobule (cluster size = 5,939 mm³, peak MNI: [−69, −24, 34.2], peak z = 4.89; supplementary Table S4). A second, smaller cluster was located in the left inferior frontal gyrus (pars opercularis), extending into the precentral gyrus (cluster size = 267 mm³, peak MNI: [−69, −3, 34.2], peak z = 3.06; supplementary Table S4). These results converge with the above findings showing that information that increases the small-world configuration of knowledge is associated with greater engagement within a distributed left-lateralized language-related cortical system.

Together, the replication provides robust evidence for behavioral preference and neural prioritization of information that increases the small world configuration of knowledge.

## Discussion

The results demonstrate that the brain encodes the potential for information to improve the structural organization of knowledge. Facts that increased the small-world index preferentially engaged the vmPFC, which is the core brain region that signals value^39,43^, as well as the language system (including the bilateral STG and left IFG) and multiple-demand system (including the dmPFC). Participants favored these facts: they liked them more and were interested in engaging with related information.

Across different systems, from biological to man-made, the small-world index is used as a measure of network efficiency^3,18–20,26,53^. It reflects the relative optimization of local connections that create clusters, and global connections across clusters. Knowledge has been shown to be organized as a small world network,^1,2^ and this organization has been associated with creativity^2,28,29^, semantic ability^2,26^, and is suggested to increase understanding^25,26,54^. Thus, the human preference we reveal here may be an adaptive response that promotes innovation and comprehension.

In fact, the discovery that information is intrinsically valued according to its capacity to improve knowledge organization, may help explain how humans are able to learn so much from so little. If the brain represents the relative value of information according to its potential to strengthen knowledge structure, it can allocate more resources towards information with high such value. This in turn can facilitate the creation of powerful mental models using sparse data. Viewed in this light, the algorithm the brain uses for information-seeking is vastly consequential for learning and reasoning.

Past accounts of information-seeking have highlighted that curiosity is driven by the need to reduce uncertainty or fill knowledge gaps^38,55^. Other accounts have suggested that people preferentially seek information that confirm what they already know^56,57^, and are especially curious about concepts they think about often, such as the self^56^. On the surface these accounts may seem contradictory; confirmation seeking rarely accords with high uncertainty reduction. Yet, these different accounts may all be understood as special cases that increases small-world configuration. A confirmatory fact, or one about a highly activated concept, may reinforce local clustering and/or reduce path length, increasing the SWI. A fact that fills a knowledge gap may do the same and/or introduce a long-range shortcut between previously disconnected concepts, reducing average path length and raising SWI. Rather than requiring separate mechanisms for these phenomena, the small-world framework offers a single computational vocabulary in which diverse influences on information preference appear as instances of a more general optimization principle.

A key insight from the present work is that this preference is best explained by the composite small-world index rather than by its individual components. Although both increases in clustering and reductions in path length independently predicted liking and wanting, a model comparison revealed that a model combining them into a single small-world metric provided a superior fit. This indicates that the human evaluation of information is sensitive to the balance between local coherence and global integration, rather than to either property in isolation. In other words, information is valued not only if it reinforces existing categories or because it connects distant concepts, but because it achieves both. This is true even when controlling for participants’ ratings of how surprising the fact is and also for cosine similarity between the concepts conveyed. The observed preference for information that increases SWI may therefore reflect an implicit drive to optimize the trade-off between semantic integration and segregation.

Interestingly, unlike the language network, BOLD response in the vmPFC was best accounted for by the ability of information to create links between distant paths. We have recently shown that while the behavioral expressions of value (i.e., liking and wanting) are often reflective of multiple factors, the traditional reward system does not necessarily track all these factors^43^. This suggest that the hedonic response of liking is not solely dependent on what has been referred to as the traditional reward system. Here, we report another such case; people clearly prefer information that optimize clustering and the creation of links between them, and the brain encodes these factors, but not necessarily both in the vmPFC.

A strength of the current study is the replication of both the fMRI results and the behavioral results using different stimuli and knowledge networks, with the behavioral findings being replicated in both online and in-person samples. Moreover, the preference for information that increases small-world configuration was observed not only in the human brain and behaviour, but also expressed by large language models. This suggests that this preference is embedded in the human-generated text the LLMs were trained on, such that an LLM can infer this preference. It does not suggest that LLMs act on such preference themselves. To the contrary, we believe the results raise the intriguing possibility that one of the reasons humans need much less data to learn than intelligent machines may be because humans use an adaptive algorithm to select what they want to know. Developing machines that incorporate similar information prioritization strategies may reduce the enormous resources that are currently needed to train them.

It is of note that the results reported here are based on a proxy of a knowledge network built using Wikipedia, that is neither fully comprehensive nor personalized. Obviously, everyone’s network will be different and much larger than the population-level model we used here. Nevertheless, the proxy network was sufficient to detect reliable group-level effects in both brain and behavior. We thus predict that the actual effects are likely much larger. We are currently working on methods to create personalized knowledge graphs per participant, though we note that any method will only generate a limited proxy. A second limitation is that in our study the information used created only pairwise conceptual connections. In life information often involves relationships among three or more concepts simultaneously. Extending our method to capture such higher-order structural changes would be of interest.

In sum, we provide evidence that humans prefer information that improves the small-world organization of knowledge networks. This suggests that the structure of semantic knowledge is a goal the human brain optimizes. People do not simply want to know more, they want to know what would make what they already know more useful. Such optimization is achieved by preferentially engaging in information that can strengthen the structure of knowledge and allocating greater neural resources to such information. This in turn may facilitate the creation of powerful mental models that enable the extraordinary human ability to understand and act upon our world.

## Methods

### Experiment I (Initial Behavioral Study)

#### Participants

96 participants completed the task online via Prolific (https://www.prolific.co/). Three participants failed to answer all attention checks correctly, so were excluded from the analysis. Data of 93 participants were thus analyzed (aged 20–64, *M* = 37.84, *SD* = 11.47; 37% female). Participants received £9.00 per hour for their participation and gave informed consent in accordance with the MIT Committee on the Use of Human Subjects (COUHES) protocol #1905811810.

### Stimuli

Our stimulus set consisted of 100 short sentences conveying a fact. Sentences ranged from 4 to 19 words in length (M = 9.38, SD = 2.89), covering a wide range of topics such as zoology, science, history, and more (see Supplementary Material A). The sentences were curated by the researchers from various online sources (e.g., https://www.sciencefocus.com/science/fun-facts; https://www.rd.com/list/interesting-facts/ ). The primary selection criterion was that each sentence clearly conveyed a relationship between two concepts, concept A and concept B. For example, the sentence: “*The unicorn is the national animal of Scotland*” conveys the relationship between concept A = “*unicorn*” and concept B = “*Scotland*”.

### Procedure

The experiment was conducted online using PsychoPy and Pavlovia.com. Participants first completed a brief tutorial to familiarize themselves with the 5-point rating scale. During the main task, on each trial a fact was presented in the center of the screen for 7 seconds in random order. Participants than rated each sentence on how much they liked it on a scale from 1 (“didn’t like it at all”) to 5 (“liked it very much”) and how much they would like to read more sentences like it on a scale from 1 (“not interested at all”) to 5 (“very interested”). They also rated how surprised they were by the information presented on a scale from 1 (“not surprising at all”) to 5 (“very surprising”), which enabled us to control for prior knowledge. In addition, three attention checks were randomly embedded throughout the experiment, during which participants were instructed to select a specific response (e.g., ‘please press 1’).

### Creating Knowledge Network & calculating its SWI

#### Concepts included in the network

We created a model of a knowledge network. The network included 500 concepts. These consisted of the 169 concepts that appear in the list of 100 facts (note that some of the concept appear in more than one sentence), plus 331 additional concepts selected based on a corpus-level salience score computed as the product of global token frequency and inverse document frequency (IDF) in the Smart Data Analytics Wikipedia dataset (https://github.com/SmartDataAnalytics/Wikipedia_TF_IDF_Dataset). This measure combines a term’s overall prevalence in Wikipedia with its distribution across documents, such that higher values reflect words that are frequent in the corpus but relatively non-uniform across Wikipedia pages. Such scores provide a proxy for lexical salience, distinguishing more informative and content-bearing concepts from highly generic high-frequency words (e.g., “thing,” “person,” “data”). To evaluate the robustness of our results, we repeated the analyses using alternative network sizes of 250, 1000, and 2000 concepts. Across all variants, the main results of our analyses remained statistically significant and qualitatively unchanged.

#### Transforming concepts to embedding

We retrieved the embedding of each of the 500 concepts included in the network using *wikipedia-en-embeddings* dataset (https://huggingface.co/datasets/Supabase/wikipedia-en-embeddings). This dataset contains embedding vector representations of 224,482 Wikipedia entities, as extracted from OpenAI’s text-embedding-ada-002 model. These vectors represent that Wikipedia entries of a concept, which better convey the meaning of a concept in broader context than the embedding of a word alone (see also^58,59^)

We checked if the information described in the 100 sentences was already present in the corresponding Wikipedia pages. To do this, for each sentence, we examined the two Wikipedia pages corresponding to the two concepts mentioned in the sentence (e.g., the pages for “bread” and “currency”) and searched for the information conveyed in the sentence (e.g., ”bread was used as currency”). For 87 out of 100 stimuli the information did not appear in the Wikipedia pages.

#### Transforming embedding into a graph

To transform the retrieved embedding into a graph representation we first computed the pairwise cosine similarities between all 500 concepts. Note that cosine similarity of Wikipedia embeddings of concepts reflect the level of association between tow concepts, not how synonymous the concepts are. For example, the entry ‘Queen’ and the entry ‘Britian’ have high cosine similarity. This yielded a symmetric matrix *S* ∈ ℝ^500*X*500^, where each entry *S_i_*_j_ = 1 − *d_cos_*(*i*, *j*) represents the cosine similarity between concept *i* and concept *j*, and *d_cos_* is the cosine distance. Using this matrix, we constructed an undirected, weighted graph *G* = (*V*, *E*, *w*), where each node *v_i_* ∈ *V* corresponds to a concept, and an edge *e_i_*_j_ ∈ *E* exists between nodes *i* and *j* if *S_i_*_j_ ≥ *τ*, where *τ* is a predefined threshold. The threshold was set at the 80th percentile of the cosine similarity distribution. Importantly, the pattern of the main result remained unchanged when alternative thresholds (50th, 70th, or 90th percentiles) were applied. The weight of each edge *w_i_*_j_ ∈ *w* was set to the corresponding cosine similarity value *S_i_*_j_.

#### Computing SWI of the initial network

We computed the small-world index (SWI) of the resulting graph G, using the standard definition^18^:

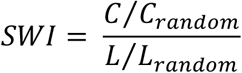

Where *C* is the clustering coefficient of the network and *L* is the average shortest path length. Because the network is weighted, we computed the clustering coefficient using the formulation proposed by.^12^ For each node *i* the weighted clustering coefficient is defined as:

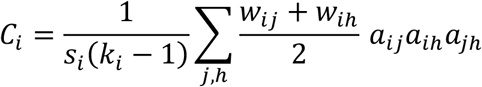

where *k_i_* is the degree (i.e., number of connections) of node *i*, *S_i_* = ∑*_j_ w_ij_* is its strength (sum of weights of incident edges), *w_i_*_j_ is the weight of the edge between nodes *i* and *j*, and *a_i_*_j_ is the element of the adjacency matrix indicating the presence of an edge between nodes *i* and *j*. The network clustering coefficient *C* is the average of *C_i_* across nodes.

The average shortest path length *L* was computed as:

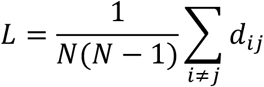

where *d_i_*_j_ is the length of the shortest path between nodes *i* and *j*, and *N* is the number of nodes in the network. In the weighted graph, path lengths were computed using the inverse of edge weights as distances.

To generate the reference random networks used to estimate *C*_random_and *L*_random_, we preserved the topology of the original graph while randomly shuffling the weights across existing edges. This procedure maintains the same number of nodes, edges, and weight distribution, while removing the original structure linking weights to specific connections.

#### Computing ΔSWI of the network following new information

To estimate the contribution of each fact to the small-world structure of the network, we artificially strengthened the connection (i.e., edge weight) between the nodes corresponding to its two key concepts. Specifically, we increased the weight *w_ab_* between the two corresponding nodes (a, b) using the formula:

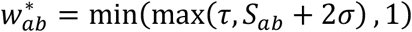

*S_ab_* is the original cosine similarity between concepts *A* and *B*, *σ* is the standard deviation of the entire similarity matrix *S*, and *τ* is the threshold value used for edge inclusion. This ensured that the new weight was at least as high as the inclusion threshold and did not exceed the maximum allowable similarity value of 1. We then recalculated the small-world index of the updated graph, and defined the change in small-worldness as:

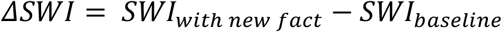

This ΔSWI value represents the change in global efficiency and modular structure of the network after receiving that piece of new information (Figure 2).

We also tested alternative methods for strengthening the *w_ab_* weight, including adding 1 standard deviations (*σ*) instead of 2*σ*, as well as directly assigning the maximum value of *S*. All variants produced qualitatively similar results, and the statistical significance of our findings remained unchanged.

#### Cosine similarity of concept A and B

During our preliminary exploratory analyses, we observed a positive correlation between ΔSWI and the cosine similarity of the vectors representing Concept A and Concept B. In light of this relationship, cosine similarity was included as a covariate in all subsequent analyses to ensure that the effects attributed to ΔSWI reflect variance beyond that explained by the semantic similarity between the concepts themselves.

### LLM experiment

The same 100 facts used in the human behavioral and fMRI studies were presented to the model OpenAI’s GPT-4o-mini via the Open AI’s Python API. For each sentence, two ratings were obtained in separate calls to ensure that no contextual memory carried over between trials. The first rating measured liking and was elicited with the prompt:

> *Here is a “fun fact” sentence containing some information:*
>
> *Sentence: {sentence}*
>
> *Rate how much you liked this sentence.*
>
> *Use the scale from “1” (did not like it at all) to “5” (really liked it).*
>
> *Important: Respond with ONLY the number (1, 2, 3, 4, or 5). No words, no explanations*
>
> *— just the single digit.*
>
> *Please use the ENTIRE range of 1–5, do NOT stick to a single rate (e.g., 4,4,4,4 or 3,3,3,3).*

The second rating measured interest in reading more similar content and was elicited with the prompt:

> *Here is a “fun fact” sentence containing some information:*
>
> *Sentence: {sentence}*
>
> *Rate how much you would want to receive and process more information like this in the future.*
>
> *Use the scale from “1” (not interested at all) to “5” (very interested!).*
>
> *Important: Respond with ONLY the number (1, 2, 3, 4, or 5). No words, no explanations*
>
> *— just the single digit.*
>
> *Please use the ENTIRE range of 1–5, do NOT stick to a single rate (e.g., 4,4,4,4 or 3,3,3,3)*.

We simulated 50 independent “participants” by running the full sentence set 50 times, each treated as a distinct subject ID.

### Statistical analyses

Behavioral and LLM data were analyzed using linear mixed-effects models, implemented in R (version 4.2.3) using the lme4 package. Separate models were fitted for each dependent variable (liking and wanting). In each model, ΔSWI was entered as a fixed effect and as a random slope over participants, with a random intercept per participant and per item. A base model included ΔSWI as the sole predictor. A second model tested the robustness of the effect by including additional covariates as both fixed and random slopes: the cosine similarity between the two key concepts in each sentence (to control for semantic relatedness) and, for the human data, the “surprising” ratings provided by participants along with participant-level demographic variables (age, gender, and education). To assess whether the composite ΔSWI measure provided better fit than either of its components individually, we fitted parallel models replacing ΔSWI with ΔC (change in the *C*⁄*C_random_* value) and ΔL (change in *L*⁄*L_random_*), as well as a combined model entering both ΔC and ΔL simultaneously as fixed effects and random slopes. Model fit was compared using the Bayesian Information Criterion (BIC), with lower values indicating better fit.

### Experiment 2 (Behavioral replication and fMRI)

#### Participants

Thirty-Four healthy volunteers were recruited via an advertisement (aged 18–51, *M*= 25.64, *SD* = 7.57; 55.5% female). 32 participants were right-handed and two participants were left-handed. Participants did not have any history of neurological, psychiatric or serious medical diagnosis, were MRI-safe and fluent in English. The study was granted ethical approval by the MIT Committee on the Use of Human Subjects (COUHES), protocol #1905811810. All participants completed informed consent and were paid $30 (£30 for participants in the UK; see below) per hour for their participation.

#### MRI Data Acquisition

A total of 34 participants underwent MRI scanning across two sites: 19 participants were scanned at the Athinoula A. Martinos Center for Biomedical Imaging, Massachusetts Institute of Technology (MIT), Boston, USA, and 15 participants were scanned at the MRC Cognition and Brain Sciences Unit (MRC CBU), University of Cambridge, UK (approved by the MRC-CBU review board #MR25009). The acquisition protocol, pulse sequences, and scanner model were identical across both sites — all data were collected on a Siemens MAGNETOM Prisma_fit 3T scanner using a head coil (HEA/HEP elements). The scanning session began with a localizer scout scan to establish participant positioning. A high-resolution T1-weighted structural image was acquired using a 3D magnetization-prepared rapid gradient-echo (MPRAGE) sequence with the following parameters: voxel size = 1.0 × 1.0 × 1.0 mm isotropic, FoV = 256 mm, 208 sagittal slices, TR = 2300 ms, TE = 2.98 ms, TI = 900 ms, flip angle = 9°, bandwidth = 240 Hz/px, GRAPPA parallel imaging with acceleration factor of 2 (32 reference lines), and 3D distortion correction applied. AutoAlign Head > Brain was used to standardize slice positioning across participants. Functional MRI data were collected using a T2*-weighted gradient-echo echo-planar imaging (EPI) sequence sensitive to BOLD contrast. Acquisition parameters were: voxel size = 3.0 × 3.0 × 3.0 mm isotropic, FoV = 210 mm, 36 axial slices acquired in interleaved order, TR = 2230 ms, TE = 30 ms, flip angle = 90°, EPI factor = 70, echo spacing = 0.49 ms, and bandwidth = 2380 Hz/px. Fat saturation was applied. Volumes were acquired continuously and the scan was terminated manually upon task completion. To enable correction of magnetic field inhomogeneities and EPI geometric distortions, a dual-echo gradient-echo field map was acquired with matched spatial coverage (voxel size = 3.0 × 3.0 × 3.0 mm isotropic, FoV = 210 mm, 35 slices, TR = 500 ms, TE1 = 3.96 ms, TE2 = 6.42 ms, flip angle = 55°, bandwidth = 1520 Hz/px). The resulting field map was used during preprocessing to apply geometric unwarping to the functional data.

#### Fmri task

The fMRI version of the task was adapted from the behavioral experiment described above and implemented in PsychoPy (v2024.2.4). The same set of 100 facts was used, presented in a random order. Participants viewed the stimuli via a mirror mounted on the head coil and made responses using an MR-compatible 5-button response box. Prior to the main task, participants read on-screen instructions explaining the procedure and confirmed their ability to read the text clearly within the scanner environment. They then completed a brief practice session to become familiar with the 5-point response scale and the button box.

Each trial followed the structure used in the behavioral study: a sentence was presented for 5 seconds, followed by a self-paced rating phase. In the fMRI version participants were asked only a single question during scanning - “How much did you like that?”- and each trial was preceded by a jittered fixation cross (3.5, 5.5, or 7.5 seconds, in random order). The task was synchronized with scanner acquisition via trigger pulses, and wall-clock timestamps were recorded throughout. The task concluded with a final screen indicating the end of the session. Total task duration was approximately 15–20 minutes, depending on individual response times. All responses, reaction times, and timing information were recorded for later analysis.

Outside of the scanner, following the scan, participants observed all 100 facts again and rated them on the three scales: ‘How much did you like this fact?’ (from 1 ‘didn’t like it at all’ to 5 ‘liked it very much’, self-paced); ‘How much do you want to learn more facts like this?’ (from 1 ‘not interested at all’ to 5 ‘very interested’); and ‘How familiar was this information to you?’ (from 1 ‘not familiar at all’ to 5 ‘very familiar’).

#### Fmri preprocessing

Functional MRI data were preprocessed using fMRIPrep 25.1.3.^61^ T1-weighted images were first corrected for intensity non-uniformity using N4BiasFieldCorrection (ANTs 2.6.2) and skull-stripped using the ANTs brain extraction workflow with the OASIS30ANTs template. Brain tissue segmentation into grey matter, white matter, and CSF compartments was performed using FSL’s FAST, and cortical surface reconstruction was carried out with FreeSurfer 7.3.2. Structural images were spatially normalized to the MNI152NLin2009cAsym standard space via nonlinear registration using ANTs.

For functional data, a reference volume was first generated for each BOLD run, after which head motion parameters were estimated using FSL’s mcflirt. Susceptibility-induced geometric distortions were corrected using the acquired dual-echo gradient-echo fieldmap, which was aligned to the EPI reference via rigid-body registration prior to unwarping. The distortion-corrected BOLD reference was then co-registered to the T1w image using boundary-based registration (bbregister, FreeSurfer) with six degrees of freedom.

Several nuisance time series were computed for confound modeling, including framewise displacement (FD), DVARS, and anatomical and temporal CompCor (aCompCor, tCompCor) components derived from white matter and CSF signals. Temporal derivatives and quadratic terms were additionally computed for head motion estimates and global signal regressors. Volumes exceeding a threshold of 0.5 mm FD or 1.5 standardized DVARS were flagged as motion outliers and subsequently modeled as nuisance regressors in the first-level GLM. All spatial transformations — including motion correction, fieldmap unwarping, and normalization to standard space — were combined and applied in a single interpolation step to minimize signal degradation.

#### First- and second-level statistical analysis

Statistical analysis was performed using Nilearn version 0.10.4 (Python version 3.10.11). For each participant, the BOLD signal was modeled using a General Linear Model (GLM) incorporating a canonical hemodynamic response function (HRF) based on the Glover model. The ‘fact’ event was defined as a boxcar encompassing the period during which each fact was presented on the screen (5s). ΔSWI was included as the primary parametric modulator of interest. Additional parametric modulators were included to control for potential confounds at the stimulus level: familiarity ratings (collected after scanning; this is equivalent to the “surprise” measurement we collected in the behavioral task.), cosine similarity between the two concepts in each sentence, number of words in the sentence, language model–based perplexity of the sentence (extracted via OpenAI’s GPT-4o API), and the serial order in which each sentence was presented. A separate regressor for rating keypress events was also included. Motion-related and physiological confounds were modeled using nuisance regressors, including six motion parameters and their temporal derivatives, white matter and cerebrospinal fluid signals (aCompCor), framewise displacement (FD), and outlier volumes identified based on DVARS (threshold = 1.5). Temporal high-pass filtering was implemented using discrete cosine basis functions with a cutoff of 128 seconds. The BOLD signal was spatially smoothed with a 6-mm full-width at half-maximum (FWHM) Gaussian kernel. Analysis was restricted to the subject-specific gray matter mask.

A group level GLM (GLM1) estimated the average effect of ΔSWI across participants, controlling for age, gender, and education. The resulting statistical map was thresholded at p < .001 (uncorrected, voxel-wise) and subsequently subjected to cluster-level family-wise error (FWE) correction at p < .05.

To further examine the BOLD response to the potential impact of a fact on network structure - specifically average path length (L) and clustering (C), we constructed an additional first-level GLM (GLM2). This model was identical to the one described above, except that instead of ΔSWI, it included ΔC (change in clustering coefficient) and ΔL (change in path length) as two parametric modulators. Both regressors were entered simultaneously to estimate their independent contributions to the BOLD signal while accounting for shared variance (r = 0.20). The same group-level analysis procedure described above was then applied.

#### ROI analyses

We conducted two ROI analyses to examine whether neural responses within the brain’s reward circuit and language network were parametrically modulated by ΔSWI. For the reward ROI, we focused on the ventromedial prefrontal cortex (vmPFC). To define this region, we used a meta-analytic approach via Neurosynth.org^47^ to generate a whole-brain association map for the term “reward” (N = 922 studies). Within the resulting vmPFC cluster, the peak voxel was identified at MNI coordinate [0, 44, 0] (Z = 7.31), and we defined a spherical ROI as a 10-mm-radius sphere centered on this peak coordinate.

For the language network ROI, we generated a meta-analytic association map for the term “language” using Neurosynth.org, thresholded using false discovery rate (FDR) correction (q < 0.01). Unlike the reward ROI, we did not restrict the language ROI to a sphere around a single peak coordinate, as language processing is broadly distributed across cortex. All surviving voxels were included, encompassing regions primarily in the bilateral superior temporal gyrus (STG) and left inferior frontal gyrus (IFG).

For each participant and each ROI, voxel-wise BOLD responses were estimated using the same GLMs as above. The resulting Beta estimates for ΔSWI were averaged across all voxels within each ROI to yield a single summary value per participant, and group-level inference was performed using one-sample t-tests against zero. To compare regression coefficients across predictors (e.g., ΔSWI vs. ΔC), we conducted paired-sample t-tests on the participant-level Beta estimates. Statistical comparisons between ΔL and ΔSWI/ΔC were conducted after inverting the scale of ΔL.

### Experiment III (Behavioral and FMRI replication)

To assess the replicability of our findings, we applied our analyses to another dataset from our lab by.^43^ Briefly, 31 different participants underwent fMRI scanning while viewing 60 pieces of semantic information and 60 visual art images, each presented for 4 seconds, rating their liking of each stimulus on a 1–5 scale. Following the scan, participants completed post-scan ratings of all stimuli on five value-related features as well as a second liking rating (see^43^ for full details of the experimental design and MRI acquisition parameters).

For the present replication, we conducted a post-hoc examination of the 60 semantic stimuli and identified 45 sentences that shared the same two-concept structure as in the current study - that is, sentences that explicitly link two distinct concepts (see Supplementary Material B). ΔSWI was computed for each of these sentences using the identical knowledge-network simulation described in the main analysis: the 90 concepts extracted from the 45 sentences were combined with 445 additional concepts selected based on high TF-IDF scores, yielding a network of 500 concepts in total. The study did not include a “wanting” scale, nor a surprise scale. However, there was a familiarity scale from 1 to 10 which was similar in spirit to the ‘surprise’ scale and thus used as a control. Thus, a linear mixed-effects model was conducted as in Exp 1 & 2, with ΔSWI as the predictor and cosine similarity, familiarity, and demographic variables as covariates.

For the neural analysis, we followed the exact same procedure as in Experiment 2 for both ROI and whole-brain analyses. The only exception was the definition of the language ROI. Instead of using the Neurosynth.org language mask employed in the main paper, we used participant-specific functional language localizers collected by^43^ as part of their study. This approach yields individually defined, functionally localized regions that better capture inter-subject variability in language responses compared to group-level meta-analytic masks. Language-selective functional regions of interest (fROIs) were defined following the procedure of^52^. Precomputed individual fROIs were obtained directly from^43^, where the full procedure is described in detail. In brief, participants completed a language localizer task in which intact sentences were contrasted with pronounceable non-word strings, presented in alternating blocks. For each participant, a sentences > non-words contrast was computed, and the top 10% of voxels most responsive to this contrast were selected within predefined language parcels^48^ to constitute the individual fROIs.

## Acknowledgements

We thank H. Haj-Ali and D. Lan for critical reading of the manuscript and helpful comments. TS is funded by the Welcome Trust Senior Research Fellowship (214268/Z/18/Z) and Discovery Award (554452/Z/25/Z). RT is funded by the fellowship of the Israel Council for Higher Education. The funders had no role in study design, data collection and analysis, decision to publish or preparation of the manuscript.

## Contributions

RT and TS designed the study, wrote the original draft and reviewed and edited the manuscript. RT performed the investigation, formal analysis and visualization. RT, AD and IP collected and preprocessed the FMRI data. IP, TN and AD provided additional guidance and reviewed the manuscript. TS acquired funding and supervised the project.

## Competing interests

The authors declare no competing interests. Data and Code Availability

All codes used to program the experiment and analyze the data will be available online via our lab GitHub at https://github.com/affective-brain-lab/SW_paper.git.

## Supplementary Materials and Tables

**Table S1.** Replication of the SWI effect on liking and wanting across various sizes of the knowledge network.

| Network Size | Liking |  | Wanting |  |
| --- | --- | --- | --- | --- |
| | $\beta$ | $p$ | $\beta$ | $p$ |
| 250 | 0.044 | < .001 *** | 0.032 | .002 ** |
| 500 | 0.052 | < .001 *** | 0.031 | .005 ** |
| 1000 | 0.039 | < .001 *** | 0.019 | .033** |
| 2000 | 0.034 | .001 ** | 0.017 | .05 |

**Note.** $\beta$ = standardized regression coefficient. Significance codes: \*\*\* $p < .001$ , \*\* $p < .01$ , \* $p = 0.05$

**Table S2.**
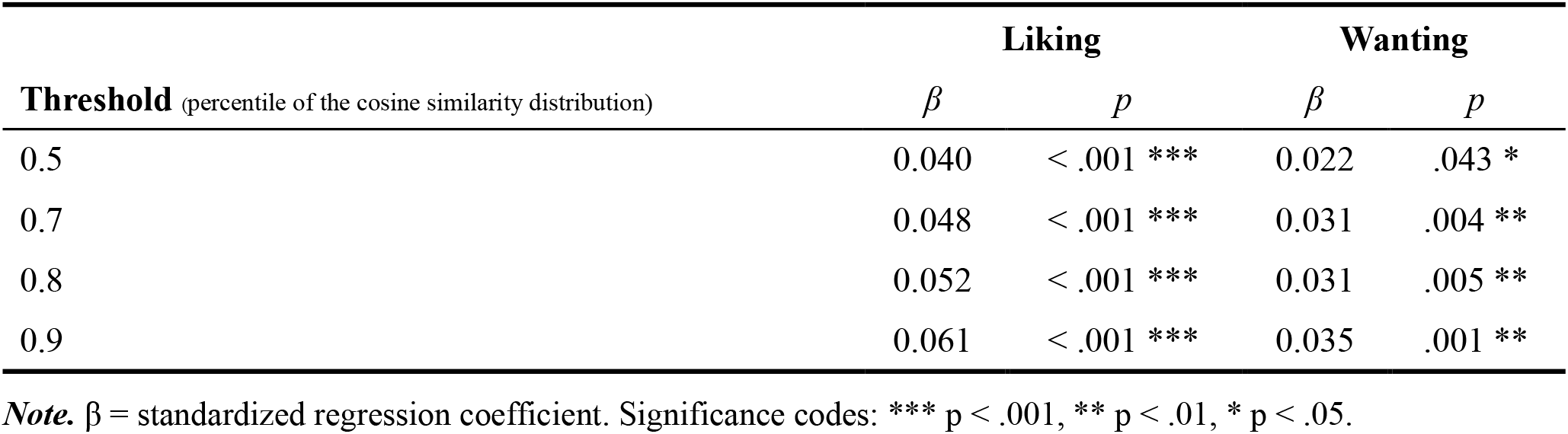
Replications of the SWI effect on liking and wanting across different knowledge networks, varying by the specific thresholds used to include edges with weights (i.e., cosine similarity) greater than or equal to the designated percentile of the distribution.

| Threshold (percentile of the cosine similarity distribution) | Liking |  | Wanting |  |
| --- | --- | --- | --- | --- |
| | $\beta$ | $p$ | $\beta$ | $p$ |
| 0.5 | 0.040 | < .001 *** | 0.022 | .043 * |
| 0.7 | 0.048 | < .001 *** | 0.031 | .004 ** |
| 0.8 | 0.052 | < .001 *** | 0.031 | .005 ** |
| 0.9 | 0.061 | < .001 *** | 0.035 | .001 ** |

**Note.** $\beta$ = standardized regression coefficient. Significance codes: \*\*\* $p < .001$ , \*\* $p < .01$ , \* $p < .05$ .

**Table S3.**
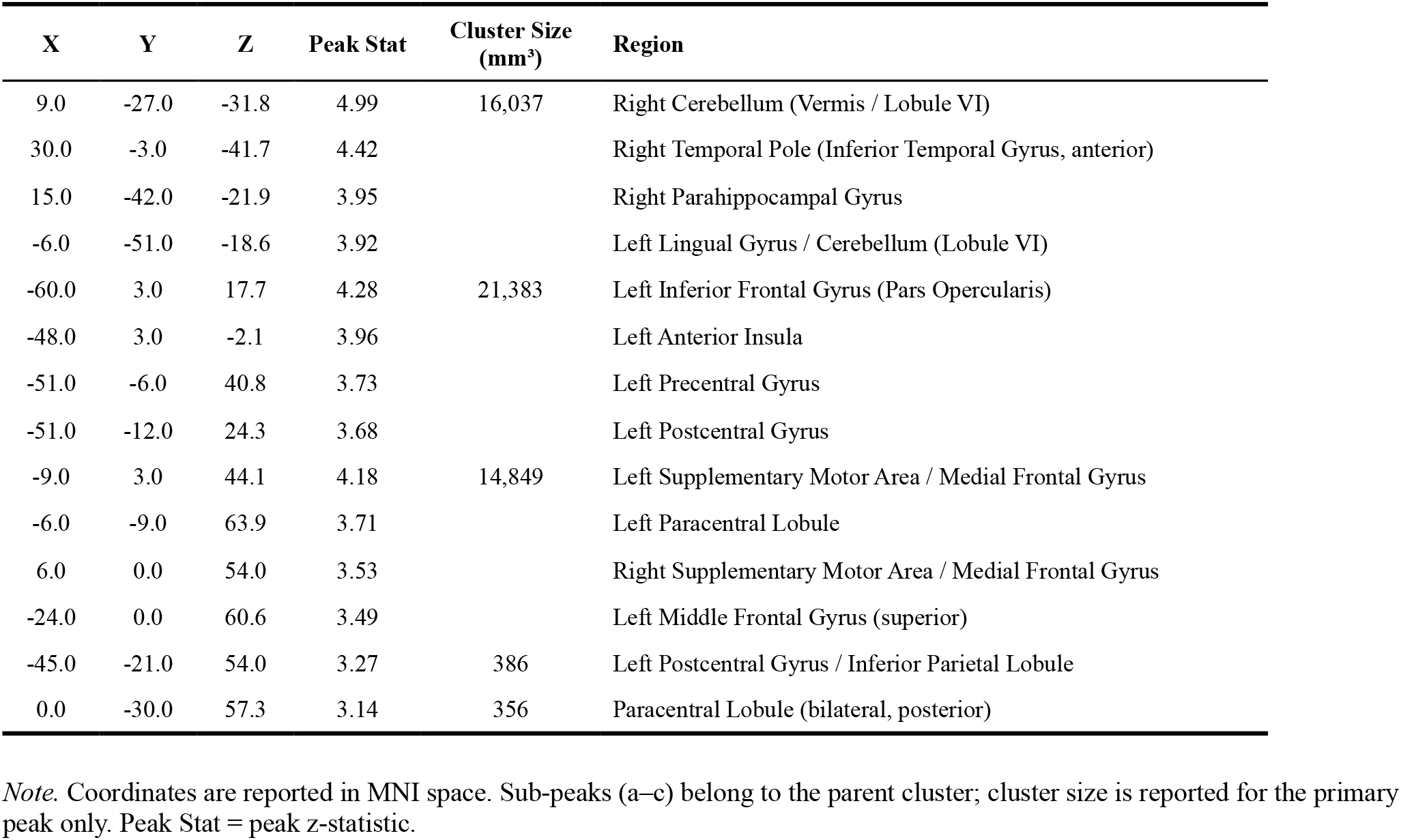
Whole-brain clusters showing significant (p<.001 voxel-wise, cluster-corrected at FWE p<.05) positive correlations with ΔSWI.

| X | Y | Z | Peak Stat | Cluster Size (mm<sup>3</sup>) | Region |
| --- | --- | --- | --- | --- | --- |
| 9.0 | -27.0 | -31.8 | 4.99 | 16,037 | Right Cerebellum (Vermis / Lobule VI) |
| 30.0 | -3.0 | -41.7 | 4.42 |  | Right Temporal Pole (Inferior Temporal Gyrus, anterior) |
| 15.0 | -42.0 | -21.9 | 3.95 |  | Right Parahippocampal Gyrus |
| -6.0 | -51.0 | -18.6 | 3.92 |  | Left Lingual Gyrus / Cerebellum (Lobule VI) |
| -60.0 | 3.0 | 17.7 | 4.28 | 21,383 | Left Inferior Frontal Gyrus (Pars Opercularis) |
| -48.0 | 3.0 | -2.1 | 3.96 |  | Left Anterior Insula |
| -51.0 | -6.0 | 40.8 | 3.73 |  | Left Precentral Gyrus |
| -51.0 | -12.0 | 24.3 | 3.68 |  | Left Postcentral Gyrus |
| -9.0 | 3.0 | 44.1 | 4.18 | 14,849 | Left Supplementary Motor Area / Medial Frontal Gyrus |
| -6.0 | -9.0 | 63.9 | 3.71 |  | Left Paracentral Lobule |
| 6.0 | 0.0 | 54.0 | 3.53 |  | Right Supplementary Motor Area / Medial Frontal Gyrus |
| -24.0 | 0.0 | 60.6 | 3.49 |  | Left Middle Frontal Gyrus (superior) |
| -45.0 | -21.0 | 54.0 | 3.27 | 386 | Left Postcentral Gyrus / Inferior Parietal Lobule |
| 0.0 | -30.0 | 57.3 | 3.14 | 356 | Paracentral Lobule (bilateral, posterior) |

**Note.** Coordinates are reported in MNI space. Sub-peaks (a–c) belong to the parent cluster; cluster size is reported for the primary peak only. Peak Stat = peak z-statistic.

**Table S4.**
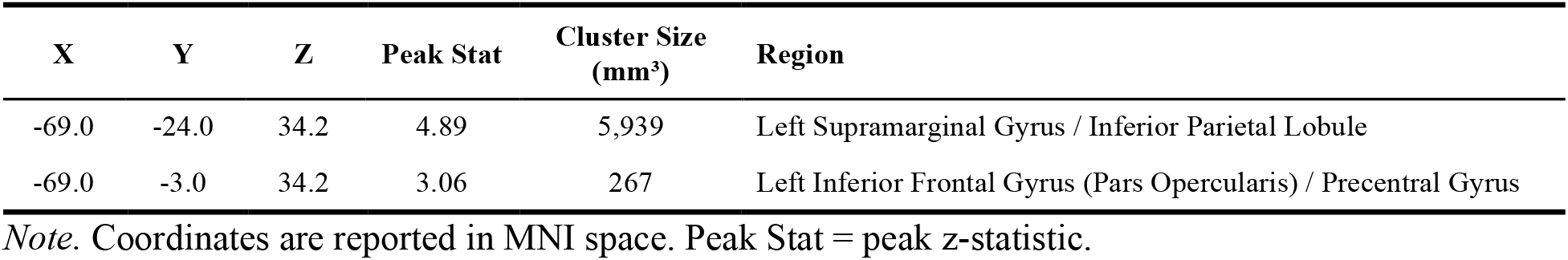
Whole-brain clusters showing significant (p<.001 voxel-wise, cluster-corrected at FWE p<.05) positive correlations with ΔSWI using Pinhorn et al’s (2026) dataset.

